# STEP inhibition prevents Aβ-mediated damage in dendritic complexity and spine density in Alzheimer’s disease.

**DOI:** 10.1101/2020.04.02.022749

**Authors:** Manavi Chatterjee, Jeemin Kwon, Jessie Benedict, Marija Kamceva, Pradeep Kurup, Paul J. Lombroso

## Abstract

Loss of dendritic spines and decline of cognitive function are hallmarks of patients with Alzheimer’s disease (AD). Previous studies have shown that AD pathophysiology involves increased expression of a central nervous system-enriched protein tyrosine phosphatase called STEP (STriatal-Enriched protein tyrosine Phosphatase). STEP opposes the development of synaptic strengthening by dephosphorylating substrates, including GluN2B, Pyk2 and ERK1/2. Genetic reduction of STEP as well as pharmacological inhibition of STEP improves cognitive function and hippocampal memory in the 3xTg AD mouse model. Here, we show that the improved cognitive function is accompanied by an increase in synaptic connectivity in cell cultures as well as in the triple transgenic AD mouse model, further highlighting the potential of STEP inhibitors as a therapeutic agent.

## 1. Introduction

Alzheimer’s disease (AD) is the most common form of dementia and leads to a progressive disruption of cognitive function. Age-dependent synapse loss and concomitant memory impairment is common in patients with Alzheimer’s (Terry et al. 1991) as well as many transgenic mouse models of AD, including the triplet transgeneic (3xTg) AD mouse model (Knobloch and Mansuy 2008). Notably, in patients with early-stage AD, dendritic spine density is reduced in the frontal cortex and hippocampal CA1 region (DeKosky and Scheff 1990; Scheff et al. 2006). The loss of spines caused by beta-amyloid (Aβ) peptide during the prodromal phase of AD has been reported to precede disintegration of neuronal networks and consequent cognitive decline (Palop and Mucke 2010; Kashyap et al. 2019). Hence, investigation of targets that could reduce spine loss is of paramount importance in understanding disease progression as well as for translational interventions.

STriatal-Enriched protein tyrosine Phosphatase (STEP) is one such target affected by Aβ accumulation in the brain. STEP is a CNS enriched phosphatase that opposes synaptic strengthening by dephosphorylating key synaptic kinases, including ERK1/2, Fyn, and Pyk2, leading to their inactivation (Venkitaramani et al. 2009; Xu et al. 2012; Li et al. 2014). STEP also dephosphorylates the GluN2B and GluA2 subunits of NMDA and AMPA receptors, leading to the internalization of these receptor complexes (Snyder et al. 2005; Zhang et al. 2008; Wu et al. 2011).

STEP is implicated in several neuropsychiatric disorders, including Alzheimer’s disease (Kurup et al. 2010; Zhang et al. 2010), Parkinson’s disease (Kurup et al. 2015), schizophrenia (Xu et al. 2018), fragile X syndrome (Chatterjee et al. 2018), as well as age-related memory decline (Castonguay et al. 2018). In AD, STEP levels are elevated in post-mortem human patients and also in several AD mouse models: Tg2576 (Kurup et al. 2010), J20 (Chin et al. 2005), APP/PS1 (Zhang et al. 2013) and 3xTG mice (Zhang et al. 2010). The accumulation of beta-amyloid (Aβ) in AD leads to the blockade of the ubiquitin-proteosome system (UPS) and activation of α7 nicotinic receptors, both processes that eventually lead to increased levels of activated STEP (Kurup et al. 2010; Zhang et al. 2013). In 3xTg-AD mice, STEP activity is significantly elevated after six months of age, which coincides with the start of memory deficits (Zhang et al. 2010).

Genetic reduction of STEP reversed the cognitive deficits experienced by 6-12 month-old 3xTg AD mouse models, thus validating STEP as a target for drug discovery (Zhang et al. 2010). The discovery and administration of a potent and specific STEP inhibitor, 8-(trifluoromethyl)-1,2,3,4,5-benzopentathiepin-6-amine hydrochloride (TC-2153), to 6-month-old 3xTg AD mice reversed the cognitive deficits in these mice (Xu et al. 2014).

As recent studies point to the involvement of STEP in spine dynamics (Cho et al. 2013b; Ng et al. 2014), we tested the hypothesis that inhibition of STEP would rescue the loss of spines in neuronal cultures and in 6-month-old 3xTg mice.

## 2. Experimental Procedures

### 2.1 Animals

All protocols were approved by the Yale University Institutional Animal Care and Use Committee and strictly adhered to the NIH Guide for the Care and Use of Laboratory Animals. C57Bl/6 and 3xTg mice on C57BL/6/SV129 background were used for the experiments. Animals were housed in groups of 2–5 in standard vented-rack cages in a 12:12 h light:dark cycle with food and water available ad libitum.

### 2.2 7PA2-conditioned medium preparation

7PA2 cells-conditioned medium (7PA2-CM; Aβ-enriched) was prepared as described previously with minor modifications (Walsh et al. 2002). The 7PA2 cells excrete Aβ oligomers, primarily dimers and trimers into the CM. Briefly, 7PA2 and untransfected Chinese hamster ovary (CHO) cell lines were grown to 95% confluency and conditioned in DMEM without serum for 16 h. Both CM as well as control CHO media was centrifuged (200g for 10 min) to remove debris and concentrated using a YM-3 column (Millipore) as described previously (Kurup et al. 2010). CM was aliquoted into 60 μl aliquots and stored at −20°C.

### 2.3 Neuronal cultures

Cortical cultures were grown from WT mice (postnatal day 0-2). For studying dendritic morphology and spine density, neuronal cultures were transfected with eGFP plasmid using DNA-CaCl2 method at day 3. Cultures were maintained for 18 days; after drug treatments, they were fixed with 4% paraformaldehyde. The cells were stained with anti-GFP and anti-MAP2 antibodies for analysis. Images were captured using Apotome at 20X (dendritic morphology analysis) and Leica confocal microscope 100X (spine density analysis). For dendritic morphology analysis, neuron tracing and subsequent Sholl analysis was done using Neurolucida software.

For synaptic puncta studies, neuronal cultures were grown for 18 days before addition of DMSO (0.1%) or TC-2153 at 1µM in the cell culture media for 48 hrs and was maintained during Aβ treatment. Cells were fixed using 4% paraformaldehyde for 10 min at room temperature. Cells were stained for the presynaptic marker, anti-synapsin (green) and the post synaptic marker, anti-PSD-95 (red). Images were captured using an Apotome microscope at 40X and analysis was performed using puncta analyzer software as described by (Ippolito and Eroglu 2010). Briefly, using the circular selection tool in the ImageJ menu to select a region radially around the soma of interest as region of interest (ROI). Both red and green channel images are thresholded and a rolling ball radius of 50 and minimum puncta size of 4 is selected. The results from the software provides quantification of red, green and colocalized puncta. Analysis was performed by researchers blinded to the experimental conditions.

**Table 1:** Materials used in the study.

| **Table 1: Materials used in the study** |  |  |
| --- | --- | --- |
| **Cat No.** | **Antibody/Drug** | **Company** |
| ab13552 | PSD-95 | Abcam |
| ab254349 | Synapsin | Abcam |
| sc-9996 | GFP | Santa cruz |
| sc-74421 | MAP2 | Santa cruz |
| SML1299 | TC-2153 | Sigma |

For spine density analysis, spines were counted from dendrites with minimum 30um length. Using Image J line tool, the spines were manually traced by drawing a line from the base of the spine to the head. The total no of spines was divided by the total dendrite length to obtain density per um value.

### 2.4 Drug treatment and Golgi staining

WT and AD mice were treated with TC-2153 or vehicle (5% DMSO in saline) at a dose of 10mg/kg, i.p. for 14 days (Xu et al. 2014). Brains were perfused with 0.9% saline and incubated in Golgi-Cox (FD Neurotechnologies) impregnation solution for 14 days. Brains were then transferred to 30% sucrose for a week before slicing into 200µm thick sections. Sections were mounted on gelatin coated slides and stained as previously described (Milatovic et al. 2010). Images were captured using a Leica confocal microscope SP5 with 40X water immersion objective using the brightfield function and quantified by tracing using multipoint tool in ImageJ analysis software. Statistical analysis was done using GraphPad Prism software.

### 2.5 Statistical analysis

All data are represented as mean ± SEM. Data from each culture was averaged and then mean values from each cultures were plotted in the graph. Statistical differences were determined using two-way ANOVA with disease and drug treatment as variables using Graphpad Prism software. P<0.05 was considered significant. Researchers were blinded to the experimental condition.

## 3. Results

### 3.1 Aβ leads to loss of dendritic complexity in primary cortical neurons, which is rescued by STEP inhibition

We first examined the effects of exogenous application of conditioned media containing Aβ on dendritic complexity in neuronal cultures. We previously demonstrated that application of Aβ for 120 min. resulted in increased STEP levels in cortical neurons with concomitant tyrosine dephosphorylation of its substrates, including a reduction in GluN2B receptors on membrane fractions (Kurup et al. 2010). We therefore treated 18-day old cortical neurons with conditioned media from mutant CHO cells containing Aβ (7PA2-CM) and control media (CM) (Fig. 1). Neurons treated with 7PA2-CM showed significantly less dendritic complexity as analyzed by Sholl analysis. There was a significant decrease in the number of dendritic junctions or nodes (P<0.01) and ends (P<0.05) compared to CM treated neurons. Application of the STEP inhibitor, TC-2153 (1μM), has previously been shown to inhibit STEP activity in cortical cultures and restore the tyrosine phosphorylation of GluN2B receptor subunits (Tyr^1472^) (Xu et al. 2014). We treated neurons with the inhibitor or vehicle for 48 hours prior to addition of either 7PA2-CM or CM. TC-2153 was maintained in culture and cells were fixed 24 hours after Aβ application and stained with MAP2 antibody. Again, we observed that neurons stimulated with 7PA2 (Fig 1C) showed significant reduction of dendritic complexity as compared with CM (P<0.05; Figure 1A). Pretreatment with TC-2153 to Aβ -treated cells significantly minimized the loss of dendritic complexity. In addition, TC-2153 pretreated Aβ cells had a greater number of nodes (P<0.05) and ends (P<0.05) compared to vehicle-treated Aβ cells (Figure 1A-G).

**Figure 1:**
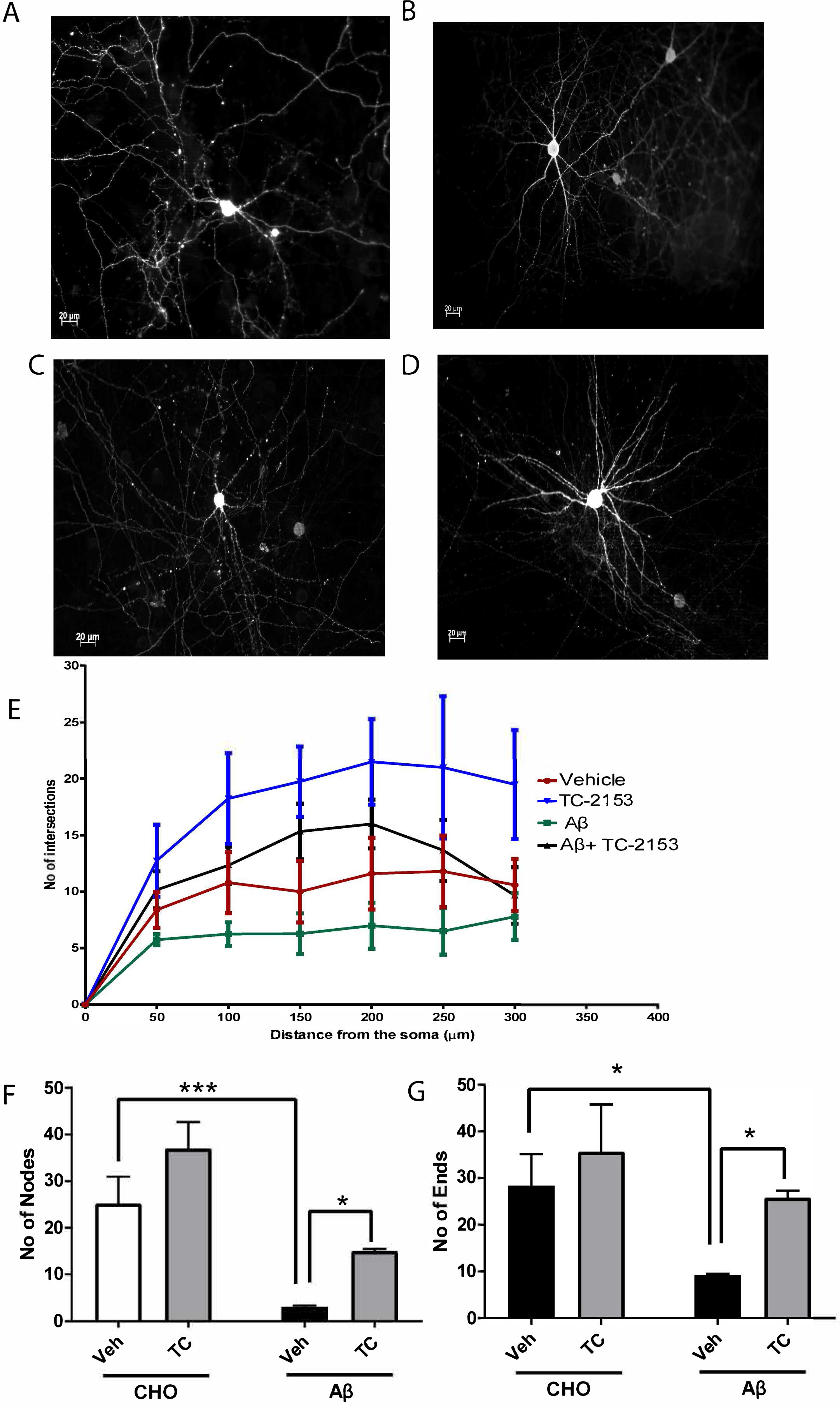
STEP inhibition prevents the loss of dendritic complexity in Aβ-treated primary cortical neurons. Primary cortical cultures (DIV-18) were treated with Vehicle (0.1% DMSO) or TC-2153 48hrs prior to addition of CHO media or conditioned media containing Aβ and immunostained with MAP2 antibody. Representative images and their respective Sholl graphs of primary cortical neuron from CHO + Veh (A); CHO+TC (B) Aβ+Veh (C) and Aβ+TC (D). Quantification of number of intersections from the soma (E). Quantification and analysis of numbers of Nodes (F) and Ends (G) across various groups. (Data from 5 neurons per culture per condition, 3-4 cultures; ***P<0.001; *P<0.05; 2-way ANOVA, Tukey post hoc analysis)

### 3.2 TC-2153 increases number of synaptic puncta in Aβ treated neurons

We next evaluated whether STEP inhibition prevented the loss of synapses in Aβ-treated neurons. We performed colocalization staining of the synaptic puncta using a pre-synaptic marker (synapsin) and a post-synaptic marker (PSD-95) and analyzed images using Puncta Analyzer (Ippolito and Eroglu 2010). Aβ-treated neurons showed significantly fewer colocalized synaptic puncta than CM-treated neurons (P<0.05) (Kono et al. 2019). Pre-treatment with TC-2153 significantly increased the colocalized synaptic puncta in Aβ-treated neurons (P<0.001) (Figure 2A-E).

**Figure 2:**
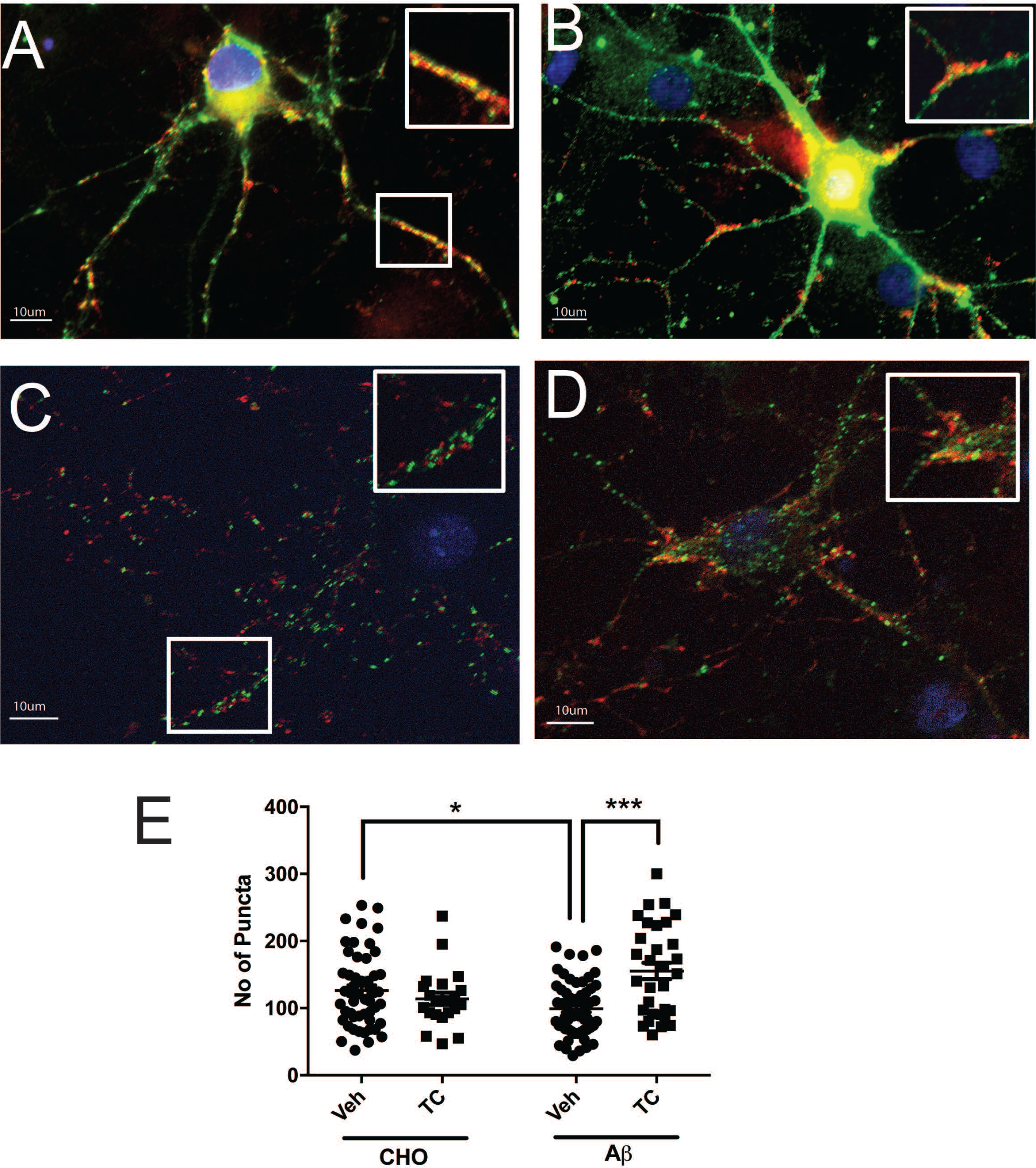
TC-2153 treatment rescues the loss of synaptic puncta in Aβ-treated neurons. WT cortical cultures were treated with TC-2153 or vehicle at DIV18, prior to addition of CHO media or conditioned media containing Aβ. Cells were stained with anti-synapsin (green) and anti-PSD-95 (red). Representative images of (A) CHO+Veh (B) CHO+TC (C) Aβ+Veh (D) Aβ+TC. Quantifications of number of colocalized puncta across all groups (E). The scale bar represents 10μm (n = 20 synapses per condition per culture, 3-4 cultures; ***P<0.001; *P<0.05; 2-way ANOVA, Tukey post hoc analysis)

### 3.3 TC-2153 attenuates spine loss in Aβ-treated neurons

Previous studies indicate that neurons exposed to Aβ show loss of spines (Calabrese et al. 2007; Shankar et al. 2007; Wei et al. 2010). Since Aβ contributes to the increase in STEP activity, we determined whether inhibition of STEP would contribute to spine loss as well. We treated transfected 3-day-old cortical neurons with pN1-EGFP and after 15 days, pretreated them with TC-2153 before adding Aβ. Consistent with previous studies, Aβ treatment resulted in a significant loss of dendritic spine density (P<0.05). However, TC-2153 pretreated cells showed no such loss of dendritic spine density (P<0.05, Figure 3E). These data indicate that TC-2153 inhibition of STEP prevents the Aβ−mediated loss of dendritic spine density.

**Figure 3:**
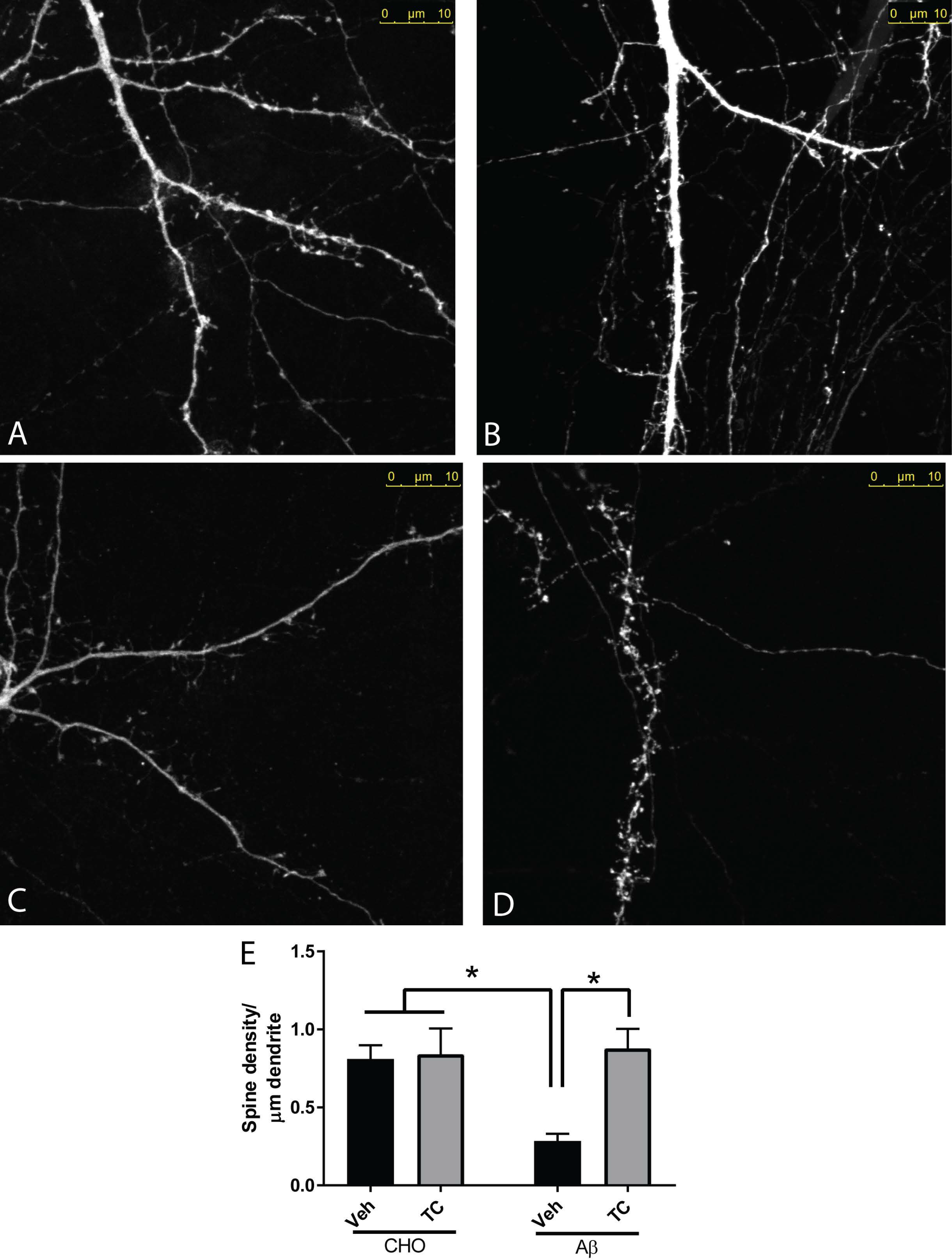
TC-2153 rescues the loss of spines in Aβ treated neurons. WT cortical cultures were transfected with eGFP on DIV-3 and treated with TC-2153 or vehicle at DIV18, prior to addition of CHO media or conditioned media containing Aβ. Cells were fixed with 4% PFA and stained with anti-GFP antibody. Representative confocal images show Representative images of (A) CHO+Veh (B) CHO+TC (C) Aβ+Veh (D) Aβ+TC are shown. Quantifications of spine density across all groups (E). Data represents Mean + SEM. (n= 5 neurons per culture per condition, 3-4 cultures; *P<0.05; 2-way ANOVA, Tukey post hoc analysis)

### 3.4 TC-2153 treatment increases dendritic spine density in 3x Tg AD mice

The increase in STEP levels in 3xTg mice is correlated with memory impairments (Kurup et al. 2010; Zhang et al. 2010). We previously showed that TC-2153 significantly rescued cognitive impairments in 6-month-old AD mice (Xu et al. 2014). To determine the effects of STEP inhibition on spine loss *in vivo*, we injected 6-month old 3xTg mice with TC-2153 (10 mg/kg, i.p.) for 14 days. Following treatment, their brains were Golgi stained and sectioned. Confocal analysis of cortical neurons showed a significant reduction of dendritic spine density in vehicetreated AD mice (P<0.01). Treatment with TC-2153 significantly rescued the dendritic spine density (P<0.05, Figure 4). These data suggest that inhibition of STEP might decrease the progression of neuronal deterioration in AD mice.

**Figure 4:**
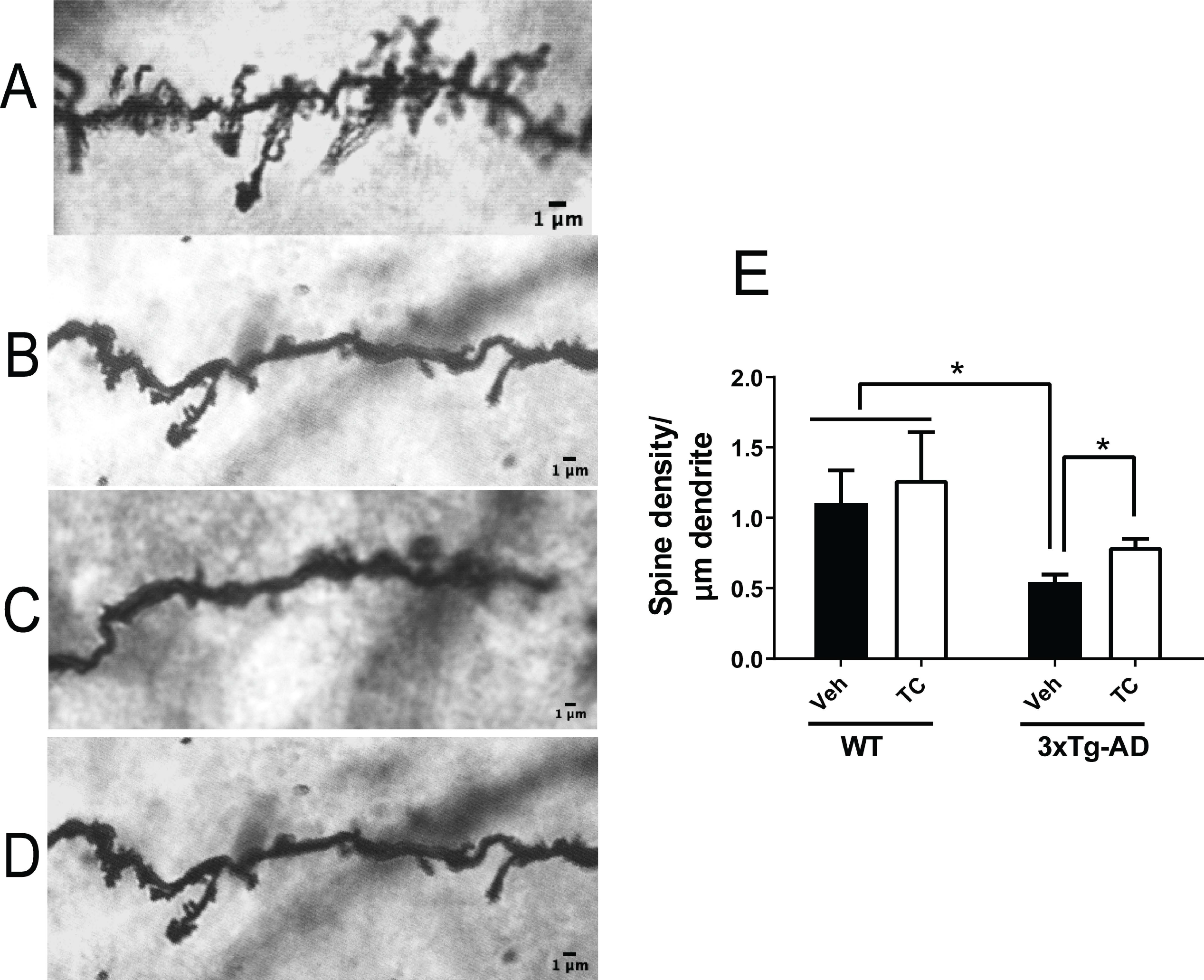
TC-2153 rescues the loss of spines in 3xTg mice. WT and 3xTg-AD mice (6-month old) were treated with TC-2153 or vehicle for 14 days. The animals were sacrificed, and the brains were collected and subjected to Golgi staining. Representative confocal images of Golgi stained sections of WT+Veh (A) WT+ TC (B) AD +Veh (C) and AD+TC (D) ;showing reduction in spine density in AD mice (C) in comparison to WT mice (A). The spine loss shows a significant improvement in mice treated with TC-2153 (E). (10 neurons per mice, n = 6 mice per group, *P < 0.05, 2-way ANOVA, Tukey post hoc analysis).

## 4. Discussion

STEP levels are elevated in Alzheimer’s disease in both post-mortem human brain samples and mouse models of Alzheimer’s disease (Kurup et al. 2010). We previously found that both genetic and pharmacological reduction of STEP activity rescued cognitive deficits in 3xTg mice (Zhang et al. 2010; Xu et al. 2014). Addtionally, STEP inhibition has been beneficial in maintaining synaptic homoeostasis in the hippocampal neurons in mouse models of fragile-X syndrome (Chatterjee et al. 2018). This is partly because activated STEP dephosphorylates NMDA and AMPA receptors and reduces their membrane expression. In this study, we provide evidence that cognitive improvements after STEP inhibition are accompanied by improvements in markers of healthy synaptic morphology, reinforcing the role of STEP in AD pathology.

STEP activity, Aβ levels, and synapse regulation are closely linked. Aβ is produced after proteolytic cleavage of amyloid precursor protein (APP). Studies have shown that Aβ oligomers reduce the density of spines in cultured neurons and transgenic mouse models (Lacor et al. 2004; Wei et al. 2010; Spires-Jones and Hyman 2014). Consistent with these structural abnormalities, neurons treated with Aβ or neurons that overexpress APP show depressed glutamatergic transmission (Ting et al. 2007). Exogenous application of Aβ induces endocytosis of NMDA receptors through a STEP-dependent pathway (Kurup et al. 2010). Since NMDA receptors are involved in the regulation of spine structure (Ultanir et al. 2007), STEP-mediated downregulation of NMDA recptors may contribute to the loss of synaptic density in AD.

Furthermore, recent studies report that F-actin (fibrillar actin), a major cytoskeletal protein that determines the shape of spines, depolymerizes to G-actin (globular actin) upon Aβ exposure, thus contributing to the collapse and decrease of spines (Kommaddi et al. 2018). We believe that SPIN90, an additional substrate of STEP, facilitates in F-actin depolymerization (Kim et al. 2007). When SPIN90 is tyrosine phosphorylated, it binds to cofilin and reduces the actin-depolymerizing activity of cofilin (Cho et al. 2013a). When SPIN90 is dephosphorylated by STEP, it leads to cofilin activation and actin depolymerization, thereby contributing to spine shrinkage (Cho et al. 2013b). Phosphorylated SPIN90 also interacts with scaffolding proteins PSD95 and Shank in the postsynaptic compartment. It is well known that Shank promotes spine maturation and enlargement (Sala et al. 2001), and that PSD95 is involved in increasing spine density and the number of synapses (El-Husseini et al. 2000). Loss of synaptic clustering with Shank and PSD95 after SPIN90 dephosphorylation by STEP affects both the size and density of dendritic spines (Cho et al. 2013a). Therefore, the inhibition of STEP using TC-2153 may prevent both the redistribution of SPIN90 in spines and possible malfunction of neural networks, resulting in improved cognition in AD mice.

## 5. Conclusion

As depicted in figure 5, our current model of AD pathophysiology suggests that Aβ inhibition of the ubiquitin degradation pathway results in increased levels of active STEP. Overactivation of STEP promotes internalization of NMDA and AMPA receptor complexes as well as dephosphorylation and translocation of SPIN90, causing spine loss after cofilin-mediated actin depolymerization. Inhibition of STEP using TC-2153 prevents this loss and rescues the pathologic synaptic morphology in conjunction with improved cognitive functioning. Long-term interventions studying TC-2153 treatment in 3xTg-AD mice, starting before the onset of memory symptoms and spine loss, may uncover the full therapeutic potential of STEP inhibitors.

**Figure 5:**
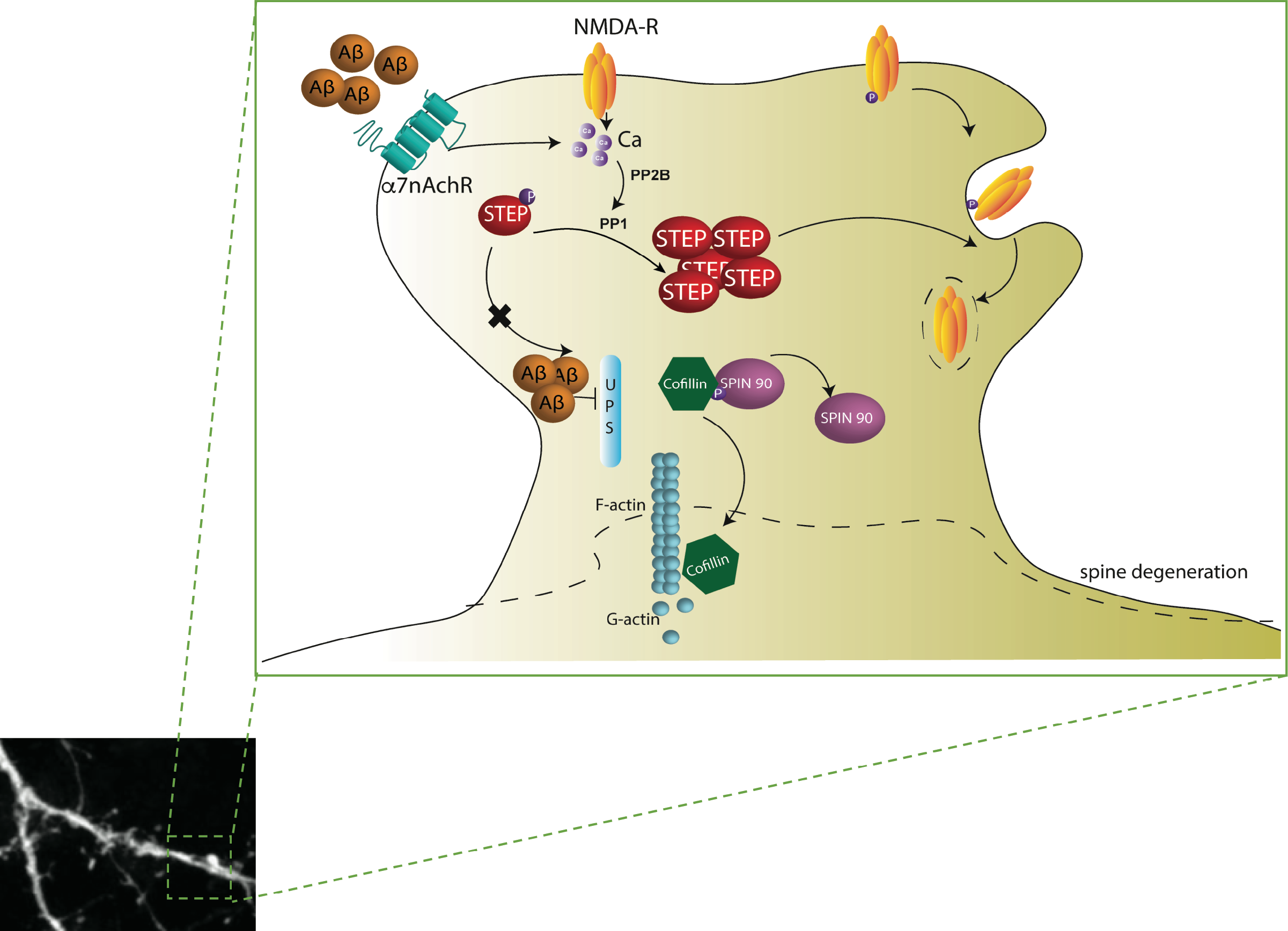
Diagrammatic representation of the current model. Aβ binding to α7nAChRs and synaptic NMDAR stimulation invokes activation of PP2B/calcineurin and PP1 to dephosphorylate STEP61, thereby increasing the affinity of STEP61 for its substrates. In a parallel pathway, Aβ inhibits the ubiquitin proteasome system (UPS) and prevents degradation of STEP61. The net result is an accumulation of active STEP protein levels in AD, which leads to increased dephosphorylation and internalization of GluN2B-containing NMDARs as well as dephosphorylation and translocation of SPIN90 to dendritic shaft. This causes release of its binding partner cofilin which severs F-actin to G-actin and thereby causes spine loss.

## Acknowledgments

We thank Ms, Clair Sulerzyski for technical assistance as well as laboratory members for helpful discussions. We are grateful to Tyler Baguley and Dr. Jonathan Ellmann of Yale Univeristy for providing TC-2153. MC and PL conceived of and designed the experiments. MC and PKK performed cell cultures and spine density studies. JK performed synaptic puncta experiments. MC, JK, JB and MK performed in vivo drug administration and Golgi staining experiments. MC and PL interpreted the data. MC, PKK and PL helped write the manuscript. All authors approved the final version of the submitted manuscript.

## Abbreviations

STEP: STriatal-Enriched protein tyrosine Phosphatase
AD: Alzheimer’s disease
3xTg: triple transgenic mouse model
CM: conditioned medium
CNS: central nervous system
ANOVA: analysis of variance
Aβ: beta amyloid
GluN2B: glutamate ionotropic receptor
NMDA type subunit 2B GluA2: glutamate ionotropic receptor
AMPA type subunit 2 Pyk2: protein tyrosine kinase 2
ERK1/2: extracellular signal-regulated kinase-1
SPIN90: SH3 protein interacting with Nck, 90 kDa
WT: wild-type

